# Structome: Exploring the structural neighbourhood of proteins

**DOI:** 10.1101/2023.02.18.529083

**Authors:** Ashar J. Malik, Chandra S. Verma, Anthony M. Poole, Jane R. Allison

**Author notes:** Correspondence to Ashar J. Malik.

## Abstract

Protein structures carry signal of common ancestry and can therefore aid in reconstructing their evolutionary histories. To expedite the structure-informed inference process, a web server, Structome, has been developed, that allows users to rapidly identify protein structures similar to a query protein and to assemble datasets useful for structure-based phylogenetics. Structome was created by clustering *∼* 94% of the structures in RCSB PDB using 90% sequence identity and representing each cluster by a centroid structure. Structure similarity between centroid proteins was calculated, and annotations from PDB, SCOP and CATH were integrated. To illustrate utility, an H3 histone was used as a query, and results show that the protein structures returned by Structome span both sequence and structural diversity of the histone fold. Additionally, the pre-computed nexus-formated distance matrix, provided by Structome, enables analysis of evolutionary relationships between proteins not identifiable using searches based on sequence similarity alone. Our results demonstrate that, beginning with a single structure, Structome can be used to rapidly generate a dataset of structural neighbours and allows deep evolutionary history of proteins to be studied. Structome is available at: https://structome.bii.a-star.edu.sg

## Introduction

The determination of protein structures has become more routine over the last two decades leading to a near-exponential growth in the number of protein structures deposited into public repositories (e.g. RCSB Protein Data Bank (PDB), rscb.org). In addition, the ability to study the dynamical behaviour of proteins may yield insights into function (1, 2, 3, 4) and aid in structure-informed drug design (5, 6, 7, 8).

The ready availability of protein structures has proven valuable for function determination (9) and evolutionary analysis of proteins (10, 11, 12, 13, 14, 15) with structure-similarity based classification databases, SCOP and CATH (16, 17), providing invaluable support in both these characterization processes. However, for novel protein structures or those not characterized by the aforementioned databases, characterization through determination of structure-based similarity remains an important step, which is further exemplified by the availability of a vast number of protein structure comparison tools e.g. TM-Align (18), MAMMOTH (19), CE (20), FATCAT (21), Dali (22, 23) and SSM (24).

Structural data is increasingly being used to assess evolutionary relationships between proteins. These relationships are conventionally explored through identifying similarities among protein sequences. However, with sequences prone to greater change, on equivalent time-scales, compared to structure (25), some evolutionary signal cannot be extracted from sequence-based analysis alone. The structure of proteins, on the other hand, being more robust to change, can retain evolutionary signal over longer time-scales and can therefore aid in the detection of homology where sequence similarity is limited (26, 27, 28, 29, 30, 31, 32, 33).

To aid in the rapid exploration of protein evolution from structure, Structome, a web-based resource for identifying structure-based relationships, is presented here.

Structome works by allowing a user to rapidly identify structures similar to an input structure. The present version was derived by taking structures from the RCSB PDB, generating clusters of similar sequences using usearch (34) and calculating structurebased similarity, using GESAMT (35) between clusters using representative (centroid) structures for each. The utility of Structome is demonstrated by using it to rapidly collect data to analyse the evolutionary history of the histone fold. Previous phylogenetic analyses of histone evolution have been at the level of individual histones (H2A, H2B, H3, H4) (36). Using Structome, we show that it is possible to rapidly generate a structural phylogeny for this fold, thus uniting individual histones plus other proteins bearing the histone fold (37) on a single phylogenetic tree.

## Methods

All data handling was carried out using the bash shell and Python v3.8.8, with numpy v1.20.1, via in-house scripts. Python was implemented using Anaconda.

### The Structome database

Protein structures and sequences were acquired from the RCSB PDB (rcsb.org). Each protein was decomposed into its constituent chains using VMD (38), followed by the implementation of a protein size (amino acid count in protein sequence) cutoff. The remaining proteins were clustered at 90% sequence identity and each cluster was represented by centroid. Each centroid was assessed and removed from Structome if it did not meet certain criterion, explained in more detail below. The final dataset of all centroid were pairwise compared using GESAMT and BLASTP to generate protein structure and sequence comparison statistics. Annotations were imported from PDB, SCOP and CATH. This data forms the Structome database, a step-by-step illustration of which is provided in Figure 1. More details surround each of these steps are given below.

**Figure 1:**
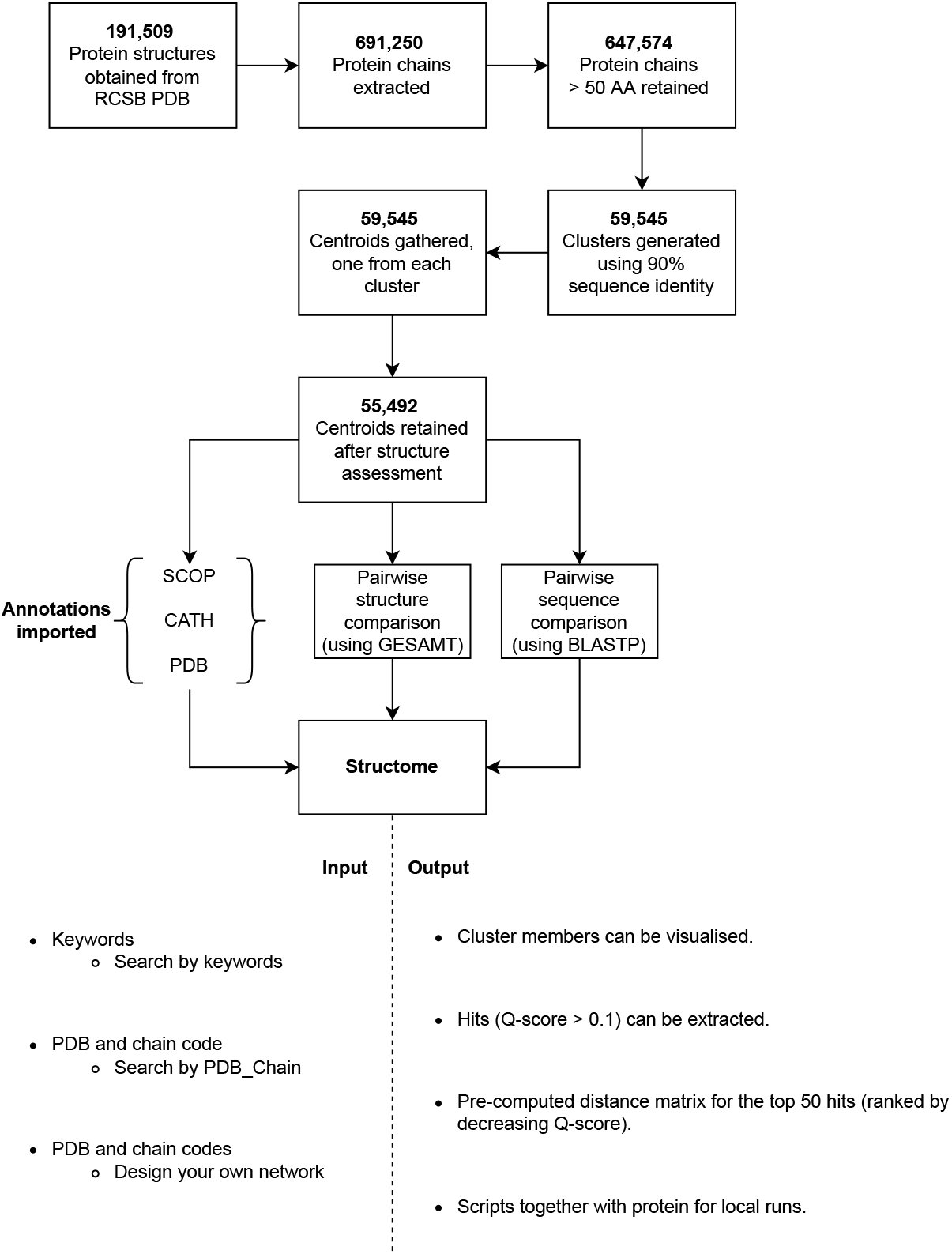
The Structome database, queries and responses. Protein structures were acquired from RCSB PDB, split into individual chains and filtered on protein sequence length. The proteins were clustered at 90% sequence identity and cluster centroid structures chosen as representatives. Protein centroids were then pairwise compared to get protein structure similarity and protein sequence similarity. This data together with annotations from SCOP, CATH and PDB form the data included in Structome. The bottom part of the diagram shows the possible queries to Structome, as keywords and PDB and chain codes, either in “Search by PDB Chain” or “Design your own network” sections. Structome, as output, allows the user to visualize the structures of all members of each cluster, for its respective centroid pick hits with Q-score *≥* 0.1, and obtain a pre-computed distance matrix, in the nexus format, for the top-50 hits, with the additional capability to download protein structures and scripts to generate distance data.

### Determination of protein size

Protein size in this context refers to the number of amino acids constituting a protein structure. This was enumerated from the sequences of all protein chains in the RCSB PDB, and only protein chains comprising 50 or more amino acids were kept for further analysis. All subsequent data on protein sizes included in this work refers to the amino acid count reported at the sequence level (i.e. Sequence length) and not the number of amino acids in the protein structure.

### Protein sequence comparison

Protein sequences were compared in two steps (see Figure 1, cluster generation and pairwise sequence comparison). In the first stage, the results of which contributed to the design of Structome, protein sequences were clustered with a cut-off of 90% sequence identity using usearch (34). The structure corresponding to the sequence of the centroid of each cluster was considered a representative of all the members in that cluster. The clustering step helped remove redundancy caused by i) multiple structures for the same protein being present ii) homo-multimeric proteins and iii) structures with point mutations.

In the latter step, using BLAST v2.8.1+ (39), a BLAST protein database was generated comprising sequences of all the centroids. Each centroid sequence was then compared to the database to provide information about sequence similarity and identity. For each comparison, the word size for BLASTP hits was changed to two amino acid to ensure a score was generated for most comparisons, accompanied by an E-value. The sequence similarity and identity data is made available to Structome users. A score of -1 was assigned to all centroid pairs where BLASTP could not find similarity across the proteins compared.

### Protein structure comparison

Secondary-structure matching (SSM) (24), has been used in earlier works (32, 33), to compare protein structures. In this work, an improved version of the same program, General Efficient Structural Alignment of Macromolecular Targets (GESAMT) (35) provided as the program gesamt in the CCP4 (Collaborative Computational Project No. 4) suite of programs (40) is used to compare protein structures.

Protein structure comparison, as with sequence comparison, was carried out in two steps (see Figure 1, centroid structure assessment and pairwise structure comparison). In the first stage, the centroid structures identified using protein sequence clustering were compared with themselves to check whether the GESAMT program was able to analyze the respective secondary structure elements in the structure. The choice of including proteins with 50 amino acids or more, is to ensure that sufficient amount of structure is present for GESAMT to work with. This check additionally ensured that all centroid protein structures could be used. If GESAMT failed to compare a protein structure with itself, another member of the cluster was chosen and subjected to the same test. This process was repeated until a suitable centroid was identified, which then replaced the original centroid. In case of a singleton cluster or if an alternate centroid could not be selected, the cluster was removed from Structome. The process was also carried out for protein structures that were provided by RCSB PDB in non-standard formats.

In the second stage of protein structure comparison, the centroid structures were compared with one another to obtain a Q-score value, which measures their degree of structural similarity. The Q-score was subtracted from one, i.e. “1 - Q-score”, to generate a distance score for the top 50 results by Q-score. This distance score is provided in the distance matrix in the nexus-formatted output file that can be downloaded from Structome for each centroid.

### SCOP and CATH classifications

For each protein structure centroid used in Structome, SCOP (41) and CATH (42) classifications were obtained, where available. Missing classifications are reflected by “N/A” in Structome.

### Protein structure visualization

Protein structure visualization is carried out using the NGL viewer (43).

### Web deployment

Structome employs the Apache server v2.4.29 with php v7.3.11 and Python v3.8.8 for server-side tasks and HTML5, CSS and JavaScript for generating the user interface.

### Illustration of usage

An analysis outside Structome was carried out, see Results (Illustration of usage). Sequence comparison was carried out between a cluster member protein and the centroid protein of the respective cluster and between two centroid proteins using BLASTP. Structural comparison was also carried out using GESAMT. The transformation matrix generated by GESAMT was used to superimpose structures using VMD.

All protein structures in the same Pfam (44) family comprising centroid protein (PF00125) were downloaded using the InterPro (45) API (see supplementary materials for Pfam data).

The centroid sequence was submitted to the HHPred online server (46, 47) (link: https://toolkit.tuebingen.mpg.de/tools/hhpred), hhsearch was used to identify protein structures that shared remote homology with the query protein using sequencebased information (see supplementary materials for HHPred results).

## Results

The motivation behind Structome was to develop a resource, which given a query protein structure could rapidly show the neighbouring protein structures based on structural similarity and, additionally, use this neighbourhood to populate a dataset which could be used to explore structure-based phylogenetic relationships. Figure 1, shows a step-by-step break down of the creation of the Structome database. This together with the web-interface and what the user may achieve is described below.

### Protein structure selection

All 191,509 PDB files (as on 22nd Sept, 2022) were downloaded from RCSB PDB. Each PDB file was split into monomers (protein chains), with each of the 691,250 chains considered an individual protein structure. Given the breadth of structures despoited in RCSB PDB, each of these chains could comprise small peptides (protein sequence length *≤* 50 amino acids), single domain or multi-domain protein.

A number of protein structures in the RCSB PDB database are short peptides (e.g, chain B, from the PDB structure 7jls, which is a tri-peptide and chain P from PDB structure 5ntn which is only a 20 amino acid long segment of a larger protein). An empirical threshold of 50 amino acids was used, and only proteins that had 50 amino acids or more were included in the Structome dataset. This size-based exclusion still retained 647,574 protein structures (i.e., *∼* 94% of protein structures in RCSB PDB) and not only removed short peptide fragments but also served the purpose of ensuring presence of sufficient structure data in the protein for GESAMT to analyze.

### Protein clustering

Several entries in the RCSB PDB comprise structures with highly similar or identical sequences. For example, some proteins are represented by a large number of structures, especially those that are commonly used to validate structure determination methods. Others are homo-multimers where each chain has an identical sequence and near identical structure, e.g. the crystal structure, PDB ID 1hv4, contains the deoxy form of hemoglobin from *Anser indicus* which is tetrameric, comprising two *α*-hemoglobin chains and two *β*-hemoglobin chains.

In other instances, structures of the same protein with single or multiple mutations are present, e.g. in the case of tumor suppressor protein p53, several mutated structures (e.g. PDB IDs: 4kvp, 4loe, 4lof and 4lo9) are present with negligible change in protein structure. To remove the resulting redundancy in the set of structures, following the size-based exclusion, the respective protein sequences were clustered with a cut-off of 90% sequence identity using usearch. The usearch program identified 59,546 clusters (reduced to 55,492 after further structure-based processing explained in the following section) based on the sequence identity cut-off. The centroid of each cluster was considered to be representative of that cluster, and subsequent sequence and structure comparison was carried out only for these centroids. This greatly reduced the redundancy in the protein structure dataset while simultaneously making it computationally tractable. The complete list of structures (after size-based filtering) can be queried to identify members of respective clusters in Structome, with the process of doing so described in some detail below.

The five largest clusters were found to have 3,631, 1,602, 1,426, 1,413 and 1,304 members, respectively, with their centroids being capsid protein (PDB ID 2gon, chain A), *β*-2 microglobulin (PDB ID 1ypz, chain B), ubiquitin-related (PDB ID 1yiw, chain A), spike glycoprotein (PDB ID 7eya, chain N) and actin (PDB ID 1d4x, chain A). The reason for the presence of the capsid protein in the top five is due to the presence of complete viral capsid structures, such as PDB ID 3j3q and 3j3y, which have 1,356 and 1,176 units of the HIV capsid protein, that is these proteins are multimers. Actin like the capsid cluster also has several multimers. The remaining three centroids are either model systems frequently used for testing methods or have important immunological roles (e.g., the spike glycoprotein), which explains their popularity as systems of interest. A large proportion of the clusters, 42%, have only one or two structures, and a further 22.53% have between three and five structures (Table 1), showing that there are many protein structures in the RCSB PDB that may be considered to be at least somewhat unrelated on a sequence level. The entire list of clusters and their members can be found in the supplementary material (Table S1).

**Table 1:**
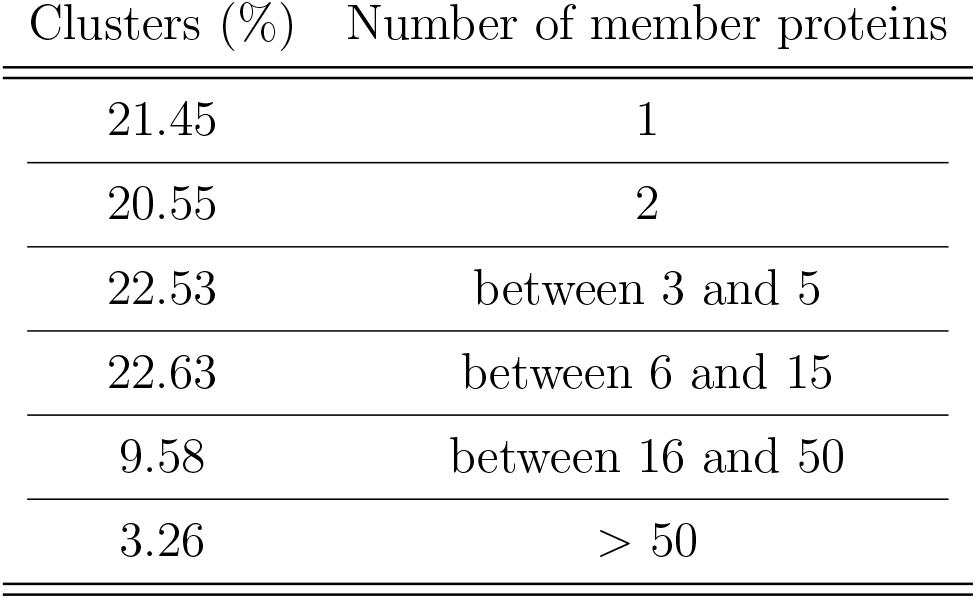
The percentage of the 55,492 clusters that contain the specified numbers of proteins.

### Protein comparison

To ensure all proteins identified by usearch as centroids could be analysed by the GESAMT structure comparison program, each centroid was obtained from the RCSB PDB, where possible, and a self-comparison was carried out. If the structure was unavailable in the standard PDB format or if the geometry of the structure was incomprehensible to GESAMT, resulting in a failed self-comparison, the centroid was replaced with another member of the cluster (see Methods). This process was repeated until either a suitable replacement was found, or there were no structures in the cluster able to be processed; in the latter case, that cluster was removed from the dataset. The numbers of clusters was subsequently reduced to 55,492 after prescreening.

The remaining centroid protein structures were pairwise compared, returning a Q-score value for each comparison. Values approaching zero indicate low structural similarity, and a value of one indicates that the two structures are identical. If the structural comparison was unable to find any structural similarity it does not report a value of zero, but instead displays a message indicating that no similarity could be detected between shared structures. In Structome, a value of -1 was assigned to such cases for ease of processing numeric data, however this was replaced with zero so that a distance value could be computed.

Every centroid when pairwise compared with other centroids generates a profile of scores ranging between zero and one. This work saw an exponential increase in the number of pairwise comparison scores below a Q-score *<* 0.1, Figure 2. This rapid change is equivalent to the increased noise in sequence-based analysis observed by Rost (48) when approaching the so-called ‘twilight zone’ of sequence similarity. Q-score values below 0.1 may therefore be considered as the structural analogy of the twilight zone. Consequently, Structome only provides comparison results that generate Q-score values *≥* 0.1, with the additional capability of selecting a higher Q-score cutoff value.

**Figure 2:**
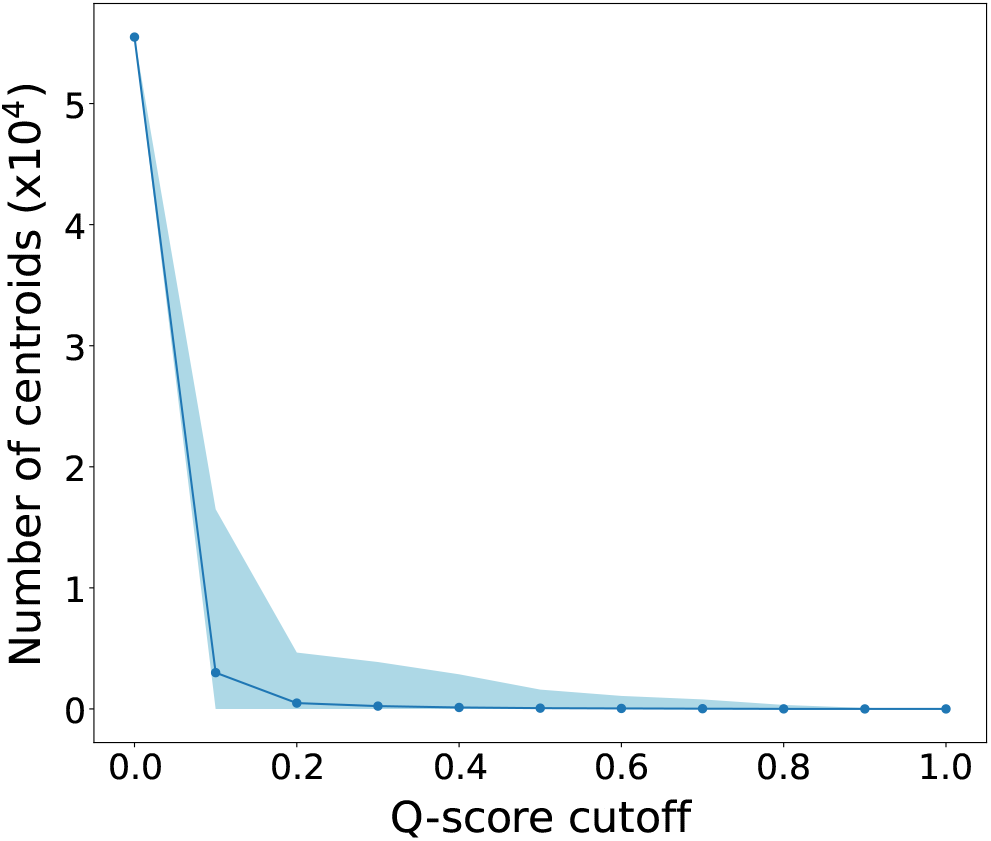
Relationship between centroid pairs and choice of a Q-score cutoff. The number of centroid pairs with Q-score values greater than a Q-score cutoff increases rapidly as the cutoff drops. The solid line represents the average number of pairs that score better than a given cutoff. The shaded region represents the amount of variation at each cutoff, for the 55,492 centroids. Detailed statistics are provided in the supplementary material (Table S1).

The sequences of each pair of centroid proteins were also compared using BLASTP to complement the structure comparison and provide an insight into any relationship between sequence and structure.

### Structome

The Structome web server provides users with a graphical web interface from which to search, organise and visualise the structure-based and sequence-based data described above. It also allows export of the structural similarity data in tabular form and as a nexus-formatted distance matrix. Below, the content and functionality of Structome is described and then illustrated with an example.

Users begin their interaction with this resource at its index page (base url : http://structome.bii.a-star.edu.sg/). This web page expects one of two kinds of input, 1) a keyword (e.g., hemoglobin) or 2) a query PDB and chain ID, (e.g., to search for chain A of PDB ID 4uuz, the search term will be 4uuz A). The chain identifier is compulsory with the latter option.

The initial response to a keyword query identifies the cluster(s) most relevant to the user input. With the use of PDB and chain ID in “Search by PDB Chain”, an exact cluster is returned. If the user query protein is classified as a centroid in Structome (e.g., the structure 1p3m A is a centroid), one row listing this structure will appear. In case of the user query not being a centroid protein structure, the centroid of the cluster to which this user query structure belongs to will be returned (e.g., the structure 4uuz A is member of the cluster represented by the centroid 1p3m A). The results are returned in the form of interactive table rows containing metadata for that Structome entry including the Structome cluster ID, centroid label, the size of the centroid protein (amino acid count), a description of the centroid as provided by RCSB PDB, its description from SCOP and CATH if present and, finally, the species in which the centroid protein is found. The last two columns of each row are action items which allow the user to either visualize all members of the cluster listed in the respective row or populate the text boxes above the preliminary results. The Q-score cutoff has a default value of 0.1 but can be increased by the user to implement a stricter cutoff. Submitting a “Cluster #”, “Centroid” and a “Q-score Cutoff” to Structome will return data describing the structural neighbourhood (within the specified Q-score cutoff) of the selected centroid structure.

The resulting page then displays a sortable table with interactive rows, but which is filled with the results of the pre-calculated protein structure and sequence comparisons between the centroid chosen in the previous step and all other centroids for which the Q-score is at least the selected cutoff value and above. The first three columns list the row number, cluster ID and the PDB ID of the centroid structure (Centroid). This is followed by four columns containing the structure comparison results: Q-score, RMSD over the aligned residues, length of the protein structure alignment (StrAln Len) and the identity between the amino acids that are structurally aligned (StrAlnSeqID). These are followed by five columns containing BLASTP sequence comparison results: number of identical residues (ID Res) from the alignment and the corresponding percentage identity (ID%),number of similar residues (Sim Res) from the alignment and the corresponding percentage similarity (Sim%) and length of the protein sequence alignment over which the percent identity and similarity are calculated (SeqAln Len). Next are columns containing the protein size and the description of the protein structure as denoted by the RCSB PDB (PDB Des). The next three columns comprise CATH and SCOP descriptions (CATH Des, SCOP Des), where available, and lastly, the species from which the protein originated.

Above the table, two histograms (left and middle, see supplementary Figure S1) summarise the protein structure comparison (Q-score) and protein sequence comparison (BLASTP sequence similarity %) scores as listed in the table. Entries with -1 are not included in the histograms. A third bar chart (right, see supplementary Figure S1), reflects a summary of the E-values generated by BLASTP comparisons. An arbitrary cutoff of E-value *≤* 0.1 was selected with the bars reflecting number of protein sequence comparisons with E-values below and above this cutoff. Given the choice of a word size of two and BLASTP returning very small sequence alignment lengths, the E-value chart helps indicate percentage of meaningful sequence comparisons.

### Data visualization and filtering

The tables provided in Structome contain useful information, but are not an intuitive way to understand the structural neighbourhood of a cluster centroid. Visualisation as a phylogenetic tree or reticulated network (see Figure 4), are much better suited to this task. To assist with visualisation, and to allow users to include protein structures not included in Structome, we have provided multiple opportunities for the data to be downloaded, as well as scripts for custom analysis.

Three buttons embedded between the charts and the table on the results page allow the user to download the data in the table. ‘Download CSV’ simply downloads all data in the table as a comma separated values format file. ‘Show Distance Data’ opens a new browser window containing a list of taxa (centroid PDB IDs) and a matrix of distance data for the 50 cluster centroids most similar to the centroid of the cluster to which the query structure belongs. The distance between two compared structures is calculated as “1 *−* Q-score” to convert the Q-score values, for which higher values correspond to more similar structures, into a measure of structural distance, or dissimilarity (30, 33). This nexus-formatted output can be downloaded for visualisation locally. Additionally, the data can be filtered by applying minimum and maximum values of the quantitative properties, or by filtering the textual descriptors for a occurrence of one or more characters (these should be entered without spaces, and are each searched for independently). Once the table contains the desired dataset, the “Download Workflow (current Table state)” button will download PDB-format files of all remaining structures in the table, along with scripts for rerunning the structure-based comparison and generating nexus-formatted distance data. At present Structome is limited by compute and therefore unable to fetch data from the database on run time for the purpose of generating a pairwise distance matrix. To circumvent this, the matrices were pre-calculated, but arbitrarily for the top 50 hits only. The feature of downloading protein structures and scripts enables the user to generate distance data for more than the pre-computed top 50 hits and ii) include their own protein structures.

Lastly, a summary page can be accessed for each cluster. This summary page can be reached via the action items in the table returned when searching Structome by keyword or PDB Chain IDs. The page can also be reached by simply clicking on on the result table rows when visualizing the hits between a centroid of choice and all other centroids in Structome. The summary page (see supplementary Figure S2) allows the user to access other members of the chosen cluster and, using the embedded graphics utility, load each of the members side by side with the cluster centroid to visually assess the degree of structural similarity. The cluster members are listed in the interactive table at the bottom of the summary page and clicking a row loads the respective structure. This list of cluster members can also be downloaded as a CSV file (Download CSV). Together, these tools allow the user to comprehensively explore the structural neighbourhood of the cluster centroid most similar to the query protein and its relationships with neighbouring proteins.

### Design your own network

Structome is currently limited by compute and therefore pre-calculated distance data in the shape of a nexus-formatted file, is only provided for the top 50 hits (by Q-score) for each centroid included in Structome. It is anticipated that users might want to analyse their own set of structures. To enable this, “Design your own network” on all web pages of Structome and “Download Workflow (current Table state)”, as discussed earlier, allows users to download PDB structures along with scripts and necessary instructions to run the analysis locally.

### Illustration of usage

Structome usage is illustrated using the structure of histone protein H3 from PDB ID 4uuz, chain A. Querying Structome with this PDB ID reveals that this structure is a member of cluster “01052”, represented by the centroid structure PDB ID 1p3m, chain A (see supplementary Figure S2).

Examination, independent of Structome, of the structures PDB ID 4uuz, chain A and PDB ID 1p3m, chain A, originating from *Drosophila melanogaster* and *Xenopus laevis*, shows that both are classified as histone H3. The two proteins also share 97% sequence identity, and have nearly identical structures (Figure 3(a)). Therefore, it is reasonable to assume that the structural neighbourhood of the centroid structure, PDB ID 1p3m, chain A, used in Structome, is likely to be analogous to that of the query structure, PDB ID 4uuz, chain A.

**Figure 3:**
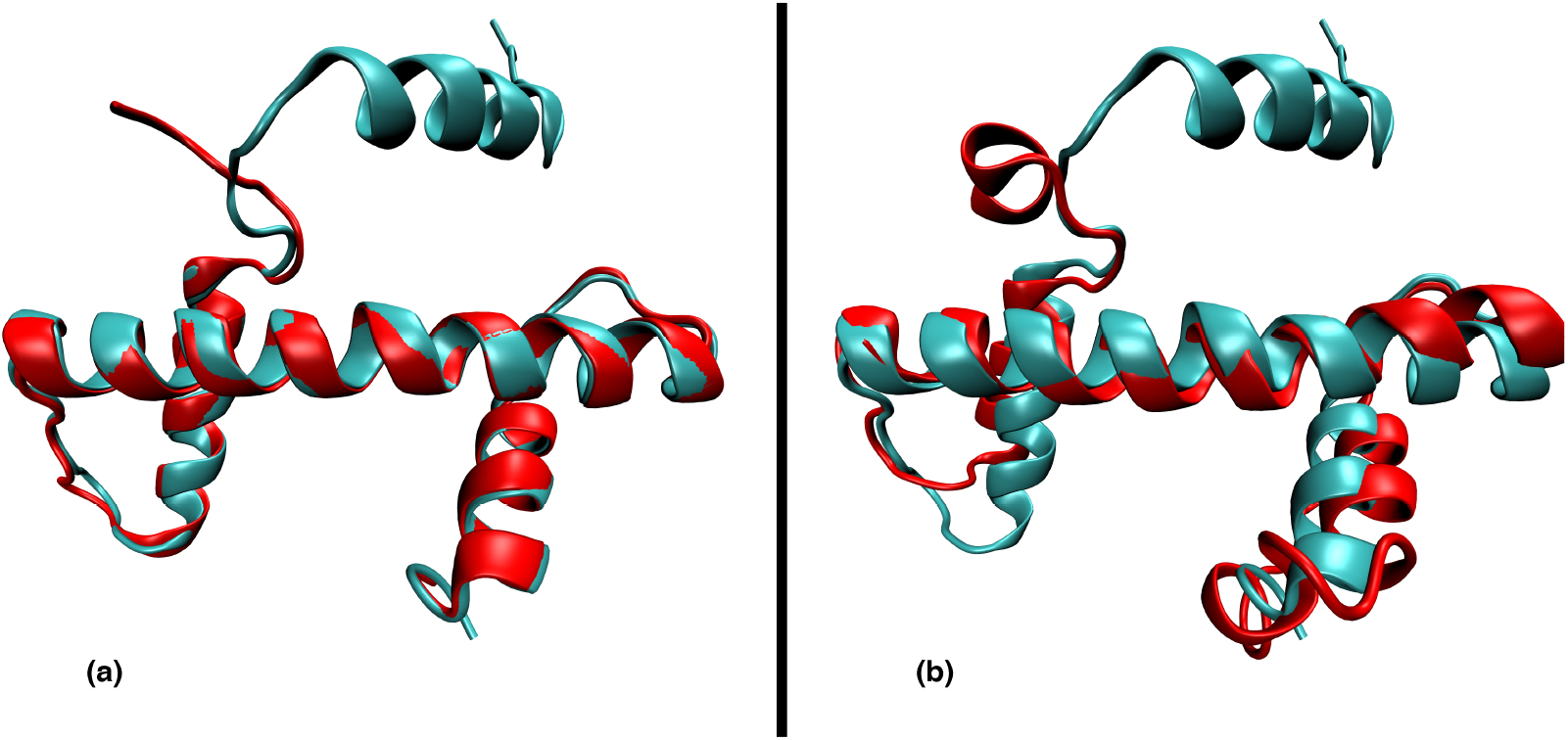
Protein structure alignment. a) The 3D protein structure alignment of PDB ID 4uuz, chain A (red) with the centroid of the cluster to which it belongs, “01052”, PDB ID 1p3m, chain A (cyan). Of their 136 and 135 amino acids, respectively, 76 could be structurally aligned. b) The protein structure alignment of the centroid of cluster “08189”, PDB ID 6m4g, chain C (red), and the centroid of cluster “01052”, PDB ID 1p3m, chain A (cyan). The Q-score for comparison of these structures is 0.409, and 72 of their 115 and 135 amino acids, respectively, can be structurally aligned; this covers the conserved histone fold(37).

The results table for centroid PDB ID 1p3m, chain A lists 5,358 centroids. These centroids have a structure similarity score Q-score *≥* 0.1, when compared to centroid PDB ID 1p3m, chain A. For these 5,358 entries, the charts (see supplementary Figure S2) show that majority of the hits have Q-scores between 0.1 and 0.3. The sequence similarity histogram shows bulk of the scores between 30% and 90%, however, given the BLASTP wordsize of two, many of the sequence similarity results are contributed by meaningless sequence alignments. To better capture this, the BLASTP E-value bar chart indicates that 5,341 (99.68%)of the hits have sequence alignments which generate E-values *>* 0.1. For instance, the sequences of centroid PDB ID 1p3m, chain A (histone-3, H3), and the centroid PDB ID 6m4g, chain C (a subtype of histone-2A, H2A), show a BLASTP alignment with 12 identical and 22 similar residues over an alignment length of 56 residues, giving a sequence similarity of 39.29%. This, alignment generates an E-value of 423,327, indicating an unreliable BLASTP sequence comparison.

Additionally, the structure-based comparison between centroid PDB ID 1p3m, chain A and PDB ID 6m4g, chain C, generates a Q-score value of 0.409, with 72 amino acids structurally aligned between them, see Figure 3(b). This and other similar entries highlight pairs that might be structurally related, however this relationship cannot be readily discerned through the alignment of their respective sequences as any comparison carried out would lie in the twlight zone of protein sequence comparison.

The pre-calculated distance matrix for the top 50 hits ranked by Q-score for the centroid PDB ID 1p3m, chain A, was obtained from Structome and visualised as a reticulated network (Figure 4). While a complete phylogenetic description is beyond the scope of this work, it is notable that the network suggests that each of the histones H2A, H2B, H3 and H4 group together with different TAFs (see partitions in Figure 4). Details regarding the origin of each taxon and its classification are provided in the supplementary material. These groupings, and additionally any phylogenetic inference obtained from it, cannot be achieved from empirical sequence-based analysis due to significant divergence at a sequence level, as demonstrated by the example earlier (see Figure 3). Additionally, centroid structures for which the sequence alignment with the centroid of choice (PDB ID 1p3m, chain A, shown in cyan in Figure 4) had E-values *≤* 0.1 are colour coded orange. This implies that starting with the H3 histone sequence (PDB ID 1p3m, chain A), BLASTP derived sequence-based similarity will only populate a dataset comprising proteins in the H3 and TAFs partition. Indeed, earlier works investigating evolutionary relationships between histones only compare histone proteins within the same families (e.g. between histone H3 from different species) and not across families (e.g. between histone H3 and H4) (36).

**Figure 4:**
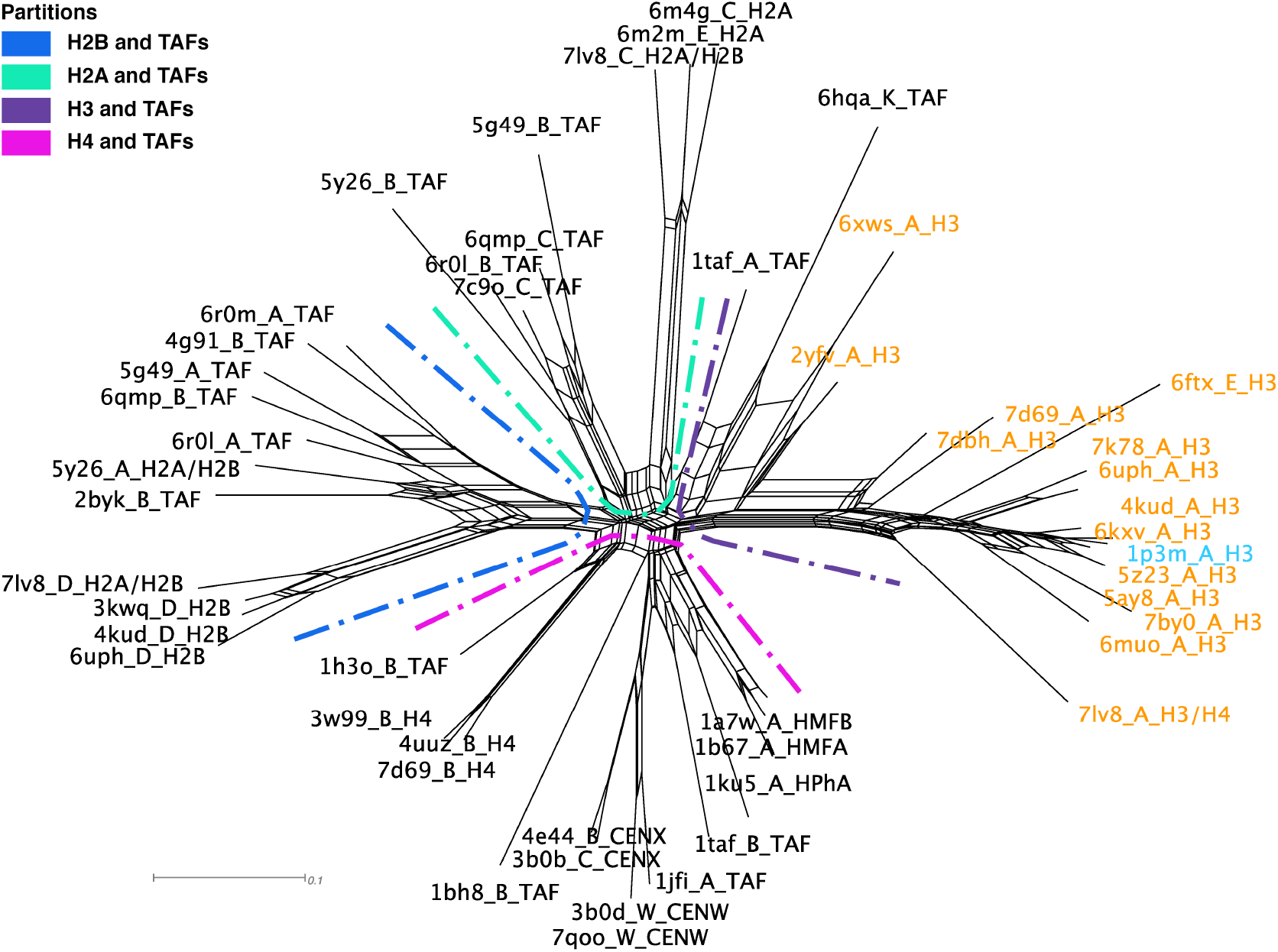
A neighbornet reticulated network of the 50 cluster centroids most structurally similar to PDB ID 1p3m, chain A (*Xenopus laevis* histone H3), shown in cyan. Each label contains the PDB ID and chain code followed by a classification obtained from RCSB PDB or literature accompanying the protein, where H2A, H2B, H3 and H4 correspond to histone proteins, TAF to TATA-binding protein-associated factors, CENX and CENW centromere proteins and HMFA, HMFB and HPhA to archaeal histones. A comprehensive list of taxa is available in the supplementary material. Departures from tree-likeness in the network indicate the existence of alternative interpretations of the data. The distance matrix was downloaded from Structome and the network was created using SplitsTree (49), with the compressed protein descriptor and partitions added during post-processing to make the network easily interpretable. The taxa in orange indicate those that are recoverable by BLASTP-based sequence similarity search resulting in E-values below 0.1. The remaining taxa either have very high E-values or are not detectable as hits by BLASTP.

A comparison with Pfam and HHPred results was also undertaken. The H3 histone (PDB ID 1p3m, chain A) is classified under the Pfam family “Core histone H2A/H2B/H3/H4” (Pfam ID PF00125). Pfam groups proteins that are homologous into families, using profile hidden Markov models, which are more sensitive than conventional BLASTP analysis. The Pfam protein family comprised 955 proteins which were separated into 3,971 protein chains (monomers). Of these 3,971 proteins, 643 proteins were not present in Structome because of size-based exclusion (e.g, PDB ID 6bhd, chain B comprising 18 amino acids, PDB ID 4i51, chain C comprising 9 amino acids etc.). Additionally 30 structures indicated by Pfam to be a member of this family returned a Q-score *<* 0.1, and hence were not considered hits. These 30 structures can generally be divided into two categories, i) where the structure is mapped to a cluster in which the centroid does not have the histone fold (e.g., PDB ID 6nzo, chain C, labelled as Ubiquitin-60S ribosomal protein L40,Histone H2A, is mapped to cluster # 01636 (using 90% sequence identity by usearch) and hence represented by centroid PDB ID 1yiw, chain A) or ii) where the structures do present a histone fold, but the proteins being compared have a significant size mismatch leading to a bias in Q-score calculations resulting in low Q-scores, explored in detail in earlier work (33). The remaining 3,298 proteins in the Pfam family, map to 52 of the 5,358 clusters returned as hits for the centroid PDB ID 1p3m, chain A. It should be noted here that Pfam family “Core histone H2A/H2B/H3/H4” groups nucleosome histones and does not exhaustively include all histones. For examples the PDB ID 6m4g, chain C, is classified under a different Pfam family (Pfam ID P0C5Z0) and has to be separately looked up.

Similar results were recovered by HHPred, another powerful method for detecting remote sequence homology, where 60 proteins were returned when querying with the sequence of PDB ID 1p3m, chain A. Of these 60, 33 proteins indicated an E-value *>* 0.1, as computed by the hhsearch program. Of the 60 proteins, three hits scored Q-score *<* 0.1 and were not considered hits by Structome, due to size difference between proteins. The remaining 57 hhsearch hits were mapped to 44 of the 5,358 clusters returned as hits for the centroid PDB ID 1p3m, chain A.

The above results demonstrate that sequence-based analysis for inferring phylogenetic relationships between i) histones themselves and ii) between histones and other DNA-binding proteins is not trivial. The ability of Structome to use a query structure to populate robust datasets, that first provide the same resolution in sequence-derived data and then capture remote signals hidden in protein structure, speaks to the value of this resource. This coupled with the capability to design networks comprising a user-selection of structures, should greatly facilitate the investigation of deep evolutionary relationships through structure-based phylogenies.

Lastly, to see all members of a particular cluster, on the centroid comparison result page, clicking the respective interactive row will provide access to cluster members. In the case of the query structure used earlier PDB ID 4uuz, chain A, clicking the top row, after sorting the results table by Q-score (descending), shows that the cluster “01052” comprises 957 protein members and is represented by the centroid PDB ID 1p3m, chain A (direct link to view just the cluster members: https://structome.bii.a-star.edu.sg/getSummary.php?Cluster=01052&PDB=1p3mA).

## Conclusion

Structome is a web server that allows users to search for protein structures that are structurally similar to a query protein. First, the centroid of the sequence similarity-based cluster to which the query structure belongs is identified, and the centroid is then used to identify other clusters for which the centroid falls within a userdefined structural similarity threshold. These centroid structures are listed along with additional information e.g, SCOP and CATH classifications. A summary page for each cluster allows users to examine other structures in the cluster. The user can export tabular results and a distance matrix for the 50 centroid structures most similar to that of the cluster to which their query structure belongs, for further analysis and visualisation as a reticulated network or phylogenetic tree. These data can be filtered prior to export, according to user-defined criteria. Lastly, the user can also export the structures and scripts to run the analysis to add value beyond what is provided by the pre-calculated distance data provided. Structome therefore provides several ways for users to identify and analyse the structural neighbourhood of any given protein structure in the RCSB PDB, with the aim of providing insight into the organisation of the protein structure landscape, in particular, with respect to the evolutionary history of proteins, through analysis of their structural relationships.

## Supporting information

supplementary statistics

supplementary statistics

## Acknowledgement

The work was supported by National Super Computing Centre (NSCC), Singpore, project number 13002493.

