## supplementary figures for "Structome: Exploring the structural neighbourhood of proteins"

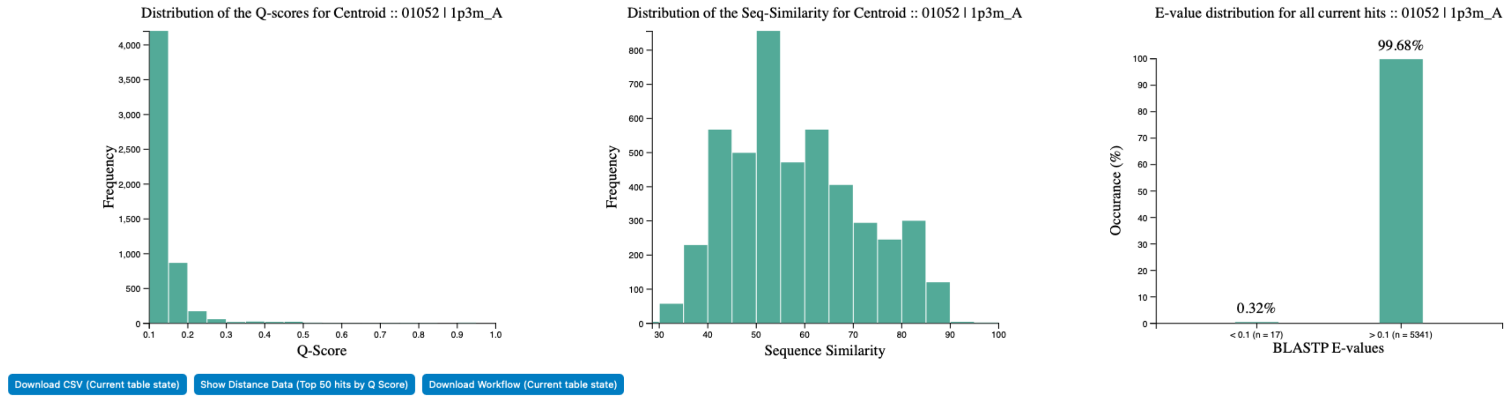


Figure S1: Summary statistics of a centroid in Structome. The charts (left and middle) show histograms of Q-score and sequence similarity of the centroid 1p3m_A compared to all other centroid in Structome that score better than the chosen cutoff of Q-score $\geq 0.1$. Given the sequence similarity calculation can return very small alignments, the bar chart on the right shows sequence comparison scoring above and below an arbitrarily chosen E-value cutoff of $\leq 0.1$. The three buttons at the bottom allow the user to download data in the table, the distance matrix for the top-50 hits and the structures and code to run locally.

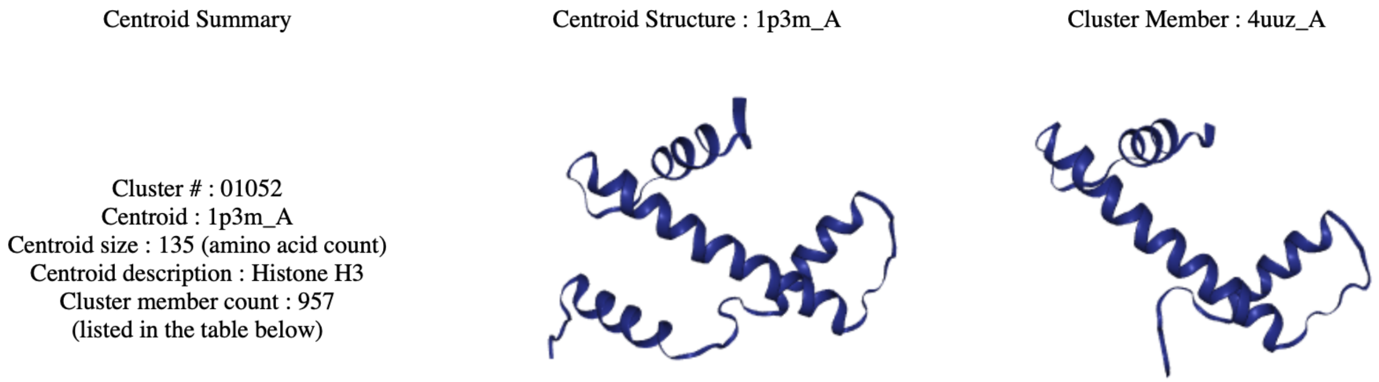


Figure S2: Centroid summary and visualization of cluster members. The left panel lists cluster number, centroid ID, centroid size in number of amino acids, centroid description as obtained from RCSB PDB, number of proteins that are member of the same cluster. The middle panel shows the protein structure of the centroid. The right panel shows all cluster members visualized one at a time using the rows in the table at the bottom of this web page to choose which cluster member to view.
